# Synapse-to-synapse plasticity variability balanced to generate input-wide constancy of transmitter release

**DOI:** 10.1101/2024.09.11.612562

**Authors:** Krisha Aghi, Ryan Schultz, Zachary L. Newman, Philipe Mendonça, Rachel Li, Dariya Bakshinska, Ehud Y. Isacoff

## Abstract

Basal synaptic strength can vary greatly between synapses formed by an individual neuron because of diverse probabilities of action potential (AP) evoked transmitter release (*Pr*). Optical quantal analysis on large numbers of identified *Drosophila* larval glutamatergic synapses shows that short-term plasticity (STP) also varies greatly between synapses made by an individual type I motor neuron (MN) onto a single body wall muscle. Synapses with high and low *P_r_*and different forms and level of STP have a random spatial distribution in the MN nerve terminal, and ones with very different properties can be located within 200 nm of one other. While synapses start off with widely diverse basal *P_r_* at low MN AP firing frequency and change *P_r_*differentially when MN firing frequency increases, the overall distribution of *P_r_* remains remarkably constant due to a balance between the numbers of synapses that facilitate and depress as well as their degree of change and basal synaptic weights. This constancy in transmitter release can ensure robustness across changing behavioral conditions.

## Introduction

Neuronal circuits must transfer information with high fidelity to ensure organismal survival. This is true in circuits where multiple similar presynaptic inputs each make only a small number of connections with multiple similar postsynaptic cells and when many synapses from the same presynaptic neuron make contact with the same postsynaptic cell (Cooper et al., 1995; Lorteije et al., 2009; Sakaba, 2018). The contributions of each synapse can vary greatly, because some synapses are reliable, with high probability of action potential (AP) evoked transmitter release (*P_r_*) and some are unreliable, with low *P_r_* (del Castillo & Katz, 1954; Dobrunz & Stevens, 1997; Hessler et al., 1993; Katz & Miledi, 1967; Mendonça et al., 2022; Newman et al., 2022).

In the face of activity-dependent plasticity change, the nervous system maintains a balanced output through the homeostatic readjustment of both synaptic strength and excitability (Davis, 2006, 2013; Eisen & Marder, 1982; Harris-Warrick & Marder, 1991; Marder & Eisen, 1984; G. Turrigiano, 2011; G. G. Turrigiano, 2008a; G. G. Turrigiano et al., 1998). However, homeostatic adjustments take hours to days for the postsynaptic compensation that regulates cell excitability and the number of neurotransmitter receptors on the cell surface, and it takes minutes for the retrograde signaling and readjustment of transmitter release of presynaptic homeostatic plasticity (PHP) (Frank, 2006). It is not clear if there are also faster homeostatic adjustments.

The *Drosophila* neuromuscular (NMJ), a model for central excitatory synapses, contains two glutamatergic motor neuron (MN) inputs that converge on common body wall muscles (Ruiz-Cañada & Budnik, 2006b, 2006a). These tonic type Ib and phasic type Is inputs differ in basal transmission, plasticity and patterns of activity during locomotion. Is MNs fire in abrupt bursts during locomotion, and their synapses release glutamate with ∼3-fold higher *P_r_* and tend to depress, whereas Ib MNs fire in ramp bursts and have lower basal *P_r_* synapses that tend to facilitate (Newman et al., 2017; Aponte-Santiago & Littleton, 2020). Both the Ib and Is axons have multiple boutons, each containing multiple active zones from where glutamate is released. When the postsynaptic response to glutamate released by both type Ib and type Is synapses is reduced (by mutation or knockdown of a subunit of the ionotropic glutamate receptor (iGluR), or by a receptor pore blocker) a retrograde signal from muscle to motor neuron triggers the compensatory increase in glutamate release of PHP (Davis, 2006, 2013; Davis & Müller, 2015; Frank et al., 2006; Nair et al., 2021; Petersen et al., 1997) but only at the type Ib input but not at the type Is input (Newman et al., 2017; Genç & Davis, 2019). Synapses of the same *Drosophila* motor axon vary greatly (over a remarkable range of 50-fold) in basal *P_r_*(Newman et al., 2022). It is not known if plasticity also varies between synapses.

Here, we ask how short-term plasticity operates across the populations of diverse Ib and Is synapses and what this means for the circuit output. We image synaptic transmission simultaneously at hundreds of identified synapses by detecting Ca^2+^ influx through the postsynaptic iGluRs with a postsynaptically targeted GCaMP, SynapGCaMP (Guerrero et al., 2004; Peled & Isacoff, 2011; Peled et al., 2014). QuaSOR, a STORM/PALM-inspired super-resolution Gaussian fitting analysis, combined with SynapGCaMP6f, provide quantal detection and spatial resolution sufficient to resolve transmission events throughout the presynaptic nerve terminals of type Ib and Is MNs, even where synapses are densely arrayed (Newman et al., 2022).

We observe a great degree of diversity in short-term plasticity, ranging by as much as 11-fold in facilitation and 13-fold in depression between synapses of similar basal *P_r_*. Short-term plasticity at Is synapses is dominated by depression, whereas Ib synapses show a broad range of plasticity. Strikingly, despite the wide range of basal *P_r_* and variability in degree and directionality of short-term plasticity between Ib synapses, the Ib motor axon as a whole displays a constancy of transmission that is preserved across a 25-fold increase in MN firing frequency. In contrast, Is synapses have a general tendency to depress. Importantly, the synapse level readjustment of transmitter release balances population transmission within seconds. We analyze the shift in balance of synaptic weight between synapses with high and low basal *Pr* and the spatial distribution of facilitating and depressing synapses and consider what the distributions mean for locomotory drive. We also examine the role of the presynaptic Gi-coupled autoreceptor, DmGluRA, which differentially inhibits Ib synapses and contributes to their *P_r_* diversity but has little to no effect on Is synapses.

## Results

### Short-term plasticity highly variant between synapses but globally constant

The *Drosophila* NMJ is innervated by two glutamatergic motor neurons, type Ib and Is, each of which contains dozens of synapses. The postsynaptic ionotropic glutamate receptors are Ca^2+^ permeable, enabling detection of synaptic transmission at the level of single synaptic quanta (vesicles) with the postsynaptically-targeted, genetically-encoded Ca^2+^ indicator SynapGCaMP6f. However, diffusion of Ca^2+^ in the postsynaptic cytoplasm makes it challenging to separate quantal events at one synapse from those arising at nearby synapses using pixel-maxima-based mapping to assign transmission site (Melom et al., 2013; Peled et al., 2014; Peled & Isacoff, 2011), meaning that events at neighboring synapses could be conflated. We recently overcame this problem with quantal synaptic optical reconstruction (QuaSOR), an analysis method that uses the fitting logic of single-molecule localization microscopy (Betzig et al., 2006; Hess et al., 2006) to enhance spatial resolution (Newman et al., 2022). Despite the fact that AP-evoked transmission can occur synchronously at neighboring synapses, responses are resolved with 2D Gaussian fits **(Figs. 1 and S1C-E)**. We QuaSOR mapped type Ib and Is postsynaptic Ca^2+^ events evoked by motor nerve stimulation at 0.2Hz (**Fig. 2A**) and aligned the transmission sites with a structural map of presynaptic active zones (AZs) obtained from Airy imaging following antibody staining for the presynaptic ELKS/CAST homolog Bruchpilot (Brp), the cytolomatrix protein that scaffolds the transmitter release apparatus (Wagh et al., 2006). From this, we determined basal synaptic transmitter release probability (*P_r_*) for each synapse as the number of (evoked events/number of stimuli) and generated a synapse *P_r_* map of ∼100-180 synapses per NMJ (**Fig. 2B**). As noted previously (Newman et al., 2022), *P_r_* varied over a wide range between synapses (**Figs. 2C and S2**).

**Figure 1.**
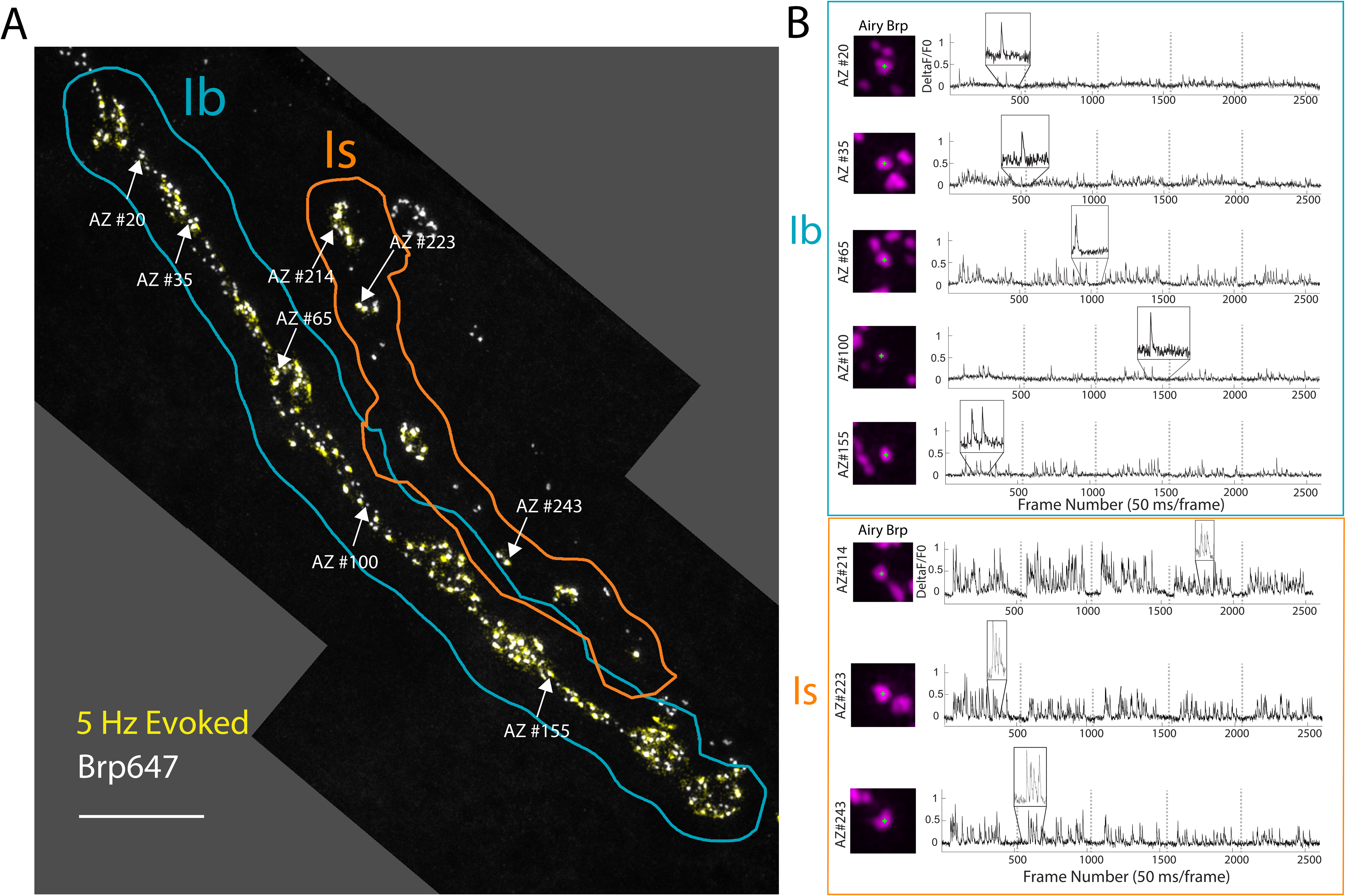
Comprehensive quantal resolution synaptic transmission imaging across active zones of Ib and Is MN inputs. A) Example probability release map showing individual release events (cyan) measured during five trains of 5 Hz evoked stimulation across Ib and Is NMJs that converge on muscle 4. Arrows point to locations of five active zones in the Ib NMJ and three active zones in the Is NMJ that differ in event densities. Scale bar: 10 µm. B) Left: Airyscan images showing Brp expression at the five identified active zones in the Ib and three identified active zones in the Is from (A). Right: ΔF/F traces tracking SynapGCaMP activity across entire evoked stimulation protocol. Grey dotted lines indicate boundaries between discrete 5 Hz trains. Inset: Traces showing quantal release events. x-axis = frame number (50 ms / frame); y-axis = normalized ΔF/F.

**Figure 2.**
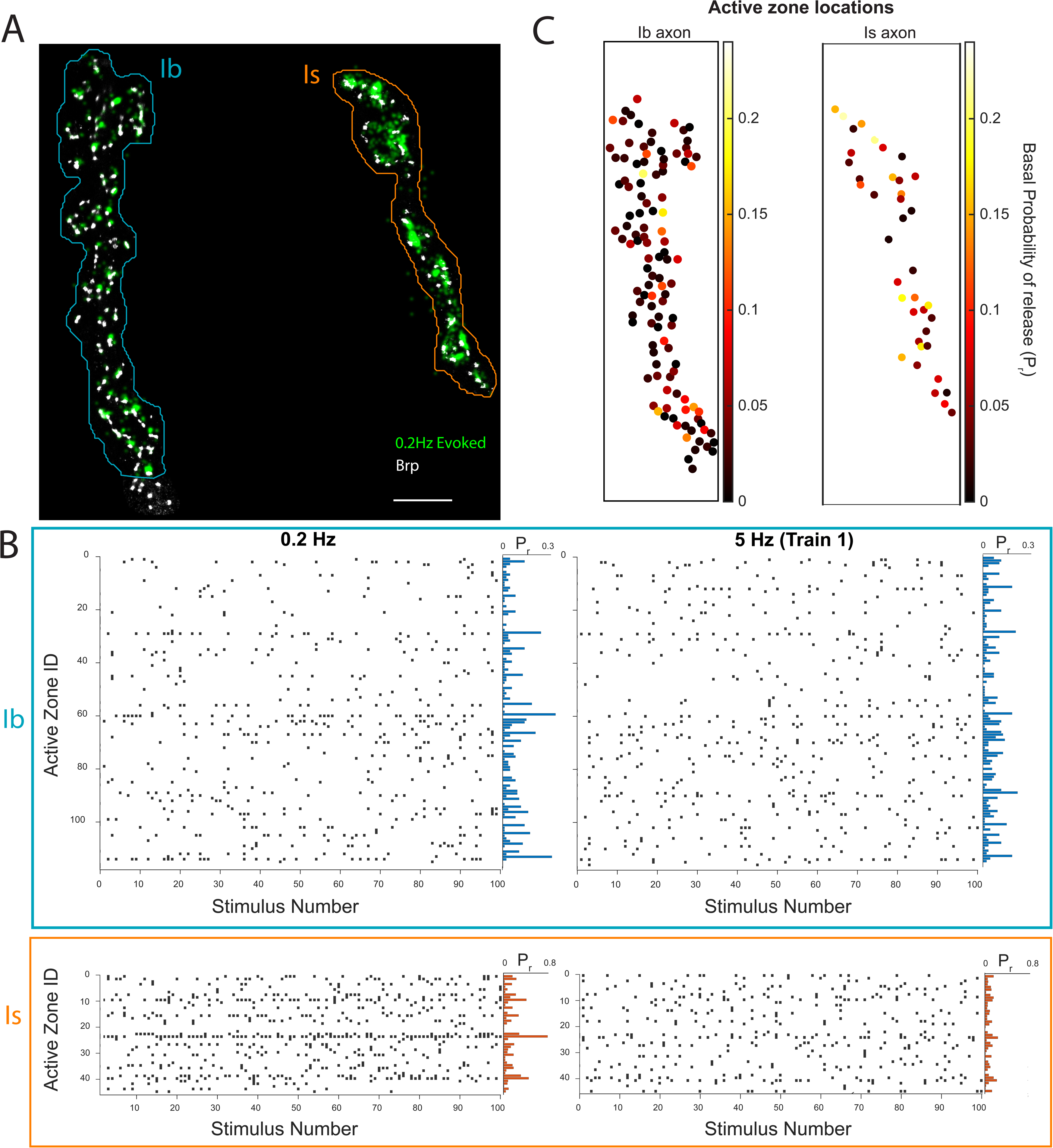
Structure-function mapping for population scale single-synapse tracking in short-term plasticity. A) Example functional Pr map from 0.2 Hz stimulation overlaid onto active zones defined by clusters of Bruchpilot (Brp) in Ib MN (outlined in light blue) and Is MN (outlined in orange) converging on muscle 4. Scale bar: 10 µm. B) Plots of each release event (vertical tick) at each identified AZ (row) in response to 100 stimuli at 0.2 Hz (left) and 5 Hz (right, showing first of the five trains). *P_r_* (average # events/stimulus) for each AZ plotted on right (Scale: Ib: 0< *P_r_* <0.3, Is: 0< *P_r_* <0.8). C) Coordinates of structurally matched AZs for the NMJs in (A) and (B). The *P_r_*of each AZ is indicated by color (scale bars on right).

Following the low frequency stimulation at 0.2 Hz, we stimulated with five trains at 5 Hz separated by rest intervals of 30 seconds and tracked transmission at each of the synapses. In the transition to 5 Hz stimulation, some synapses facilitated, some remained constant and some depressed (**Fig. 4A**). While there was a tendency toward depression for synapses with high basal *P_r_*, the degree of depression varied, and synapses with low and medium basal *P_r_* ranged between facilitation and depression, with *P_r_* changes of up to 13-fold in either direction. In the Ib, the fraction of synapses that facilitated was similar to the fraction that depressed, with the rest remaining constant, across varying levels of basal *P_r_* (facilitated: 0.41 ± 0.03; depressed: 0.40 ± 0.09; constant: 0.19 ± 0.01; N = 10 NMJs) (**Fig. 3B**). The fraction of the facilitating and depressing synapses did not differ statistically (Tukey-Kramer post-hoc comparison, *p* = 0.9618). Moreover, the degree of change between synapses that facilitated and synapses that depressed was the same (Welch’s unpaired t-test, *p* = 0.4063). For the Ib input the distribution of *P_r_* values was remarkably similar at 0.2 Hz and 5 Hz (**Fig. 4C**). Furthermore, while individual synapses varied in *P_r_* from one 5 Hz train to the next (**Fig. 4A**), the overall *P_r_* distribution for the NMJ remained constant (**Fig. S3**). In contrast, in the Is motor input the fraction of synapses that depressed was greater than the fraction that facilitated or the fraction that remained constant (facilitated: 0.28 ± 0.12; depressed: 0.65± 0.12; constant: 0.07 ± 0.03; N = 9 NMJs) (**Fig. 4B**). The degree to which synapses depressed was also greater than the degree of facilitation (Welch’s unpaired t-test, *p* = 0.001). Finally, for the Is input, the distribution of *P_r_* values shifted leftward when comparing 0.2 Hz and 5 Hz (Kolmogorov-Smirnov test, *p* < 0.0001) (**Fig. 4C**). Interestingly, facilitating, constant and depressing synapses for both Ib and Is inputs appeared to be distributed in a random fashion spatially (**Fig. 5E,F**). These results indicate that the NMJ balances the valence and magnitude of short-term plasticity across a large number of synapses in the Ib MN to maintain an overall constant output of neurotransmitter across this 25-fold range in motor neuron firing frequency. This constancy does not hold across the synapses of the Is MN.

**Figure 3.**
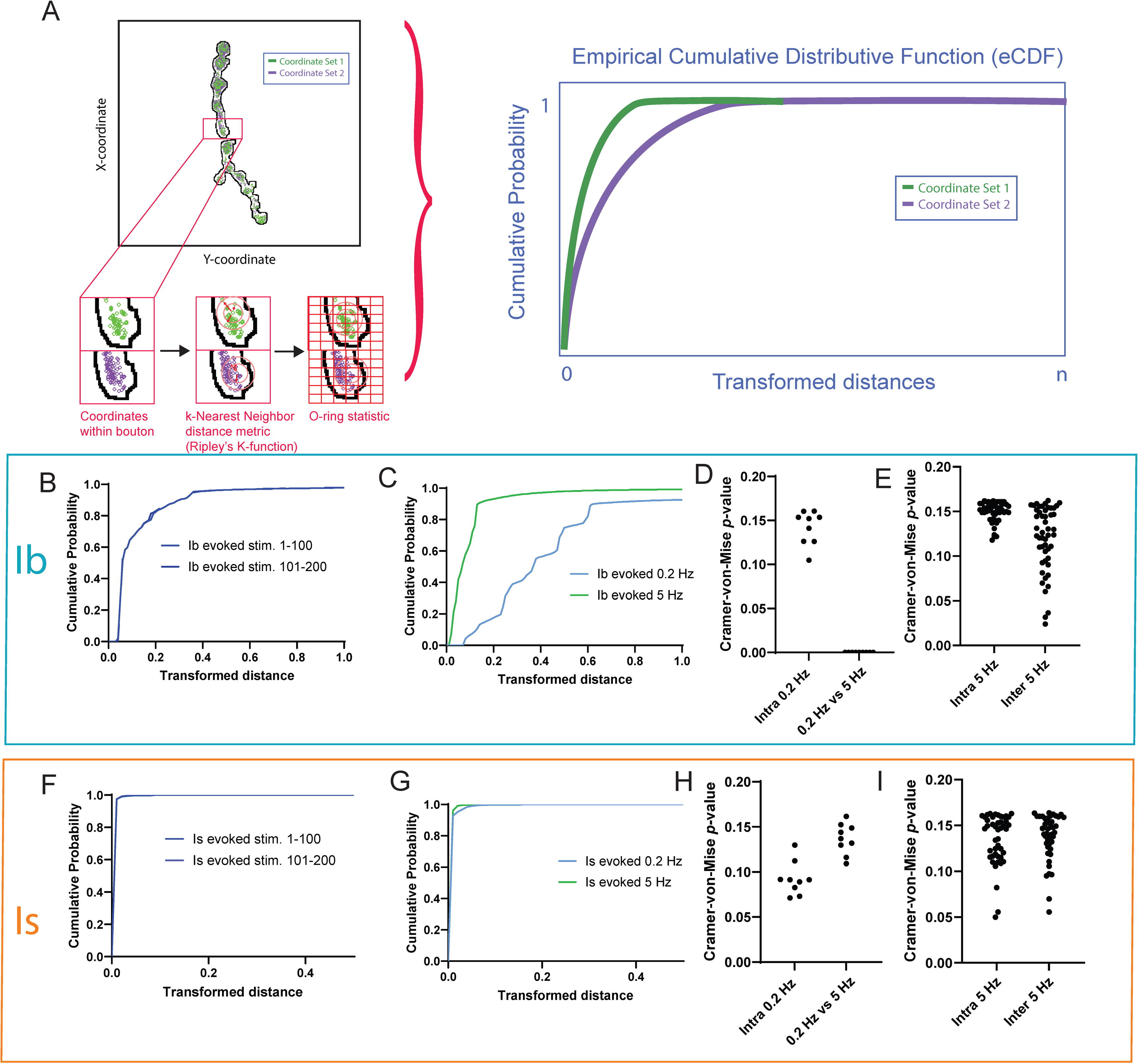
Global spatial patterns of release change with stimulus frequencies for the Ib input but not the Is input. A) Scheme depicting computation of O-ring statistic using individual evoked transmission events in example Ib axon. B) Representative transformed distance distributions for the Ib NMJ shown in (A), comparing evoked event coordinates for stimuli 1–100 (solid; *n* = 1936 events) to evoked coordinates from stimuli 101–200 (dashed; *n* = 1904 events; Cramér–von Mises *p* = 0.14). C) Ib input global decoupling of basal (0.2 Hz) and high-frequency (5 Hz) evoked transmission events. Representative transformed distance distributions for a different NMJ than in (A), comparing basal (0.2 Hz) evoked event coordinates from stimuli #1–100 (blue; *n* = 1624 events) versus event coordinates for a subsequent high frequency (5 Hz) train, stimuli #1-100, in the same NMJ (green; *n* = 1430 events; two-sample two-sided Cramér–von Mises test *p* = 0.006). D) Pooled (*n* = 9 NMJs) two-sample two-sided Cramér–von Mises test *p* values for first half stimuli (1-100) versus second half (stimuli 101-200) evoked at 0.2 Hz (left) and pooled (*n* = 10 NMJs) two-sample two-sided Cramér–von Mises test *p* values for 0.2 Hz evoked stimuli (100 stimuli) versus first 100 stimulus train at 5 Hz (right). E) Pooled two-sampled Cramer-von Mises test *p* values comparing first and second half of 100 pulse transmission events evoked at 5 Hz first (Intra 5 Hz), and comparison between different 5 Hz stimulus trains (Inter 5 Hz). F) Representative transformed distance distributions for a single Is NMJ comparing evoked event coordinates from stimuli 1–100 (solid; *n* = 2254 events) to evoked coordinates from stimuli 101– 200 (dashed; *n* = 2305 events; Cramér–von Mises *p* = 0.07). G) Different Is NMJ than shown in (F) release evoked at low (0.2 Hz; all stimuli 1–100; blue; *n* = 403 events) and high-frequency (5 Hz; all stimuli 1-100; green; *n* = 339 events) have similar transformed distance distributions (two-sample two-sided Cramér–von Mises test *p* = 0.132). H) Pooled (*n* = 9) two-sample two-sided Cramér–von Mises test *p* values for evoked first half versus evoked second half and pooled (*n* = 9) two-sample two-sided Cramér–von Mises test *p* values for 0.2 Hz evoked stimuli versus 5 Hz evoked cumulative distributions for multiple WT Is NMJs. I) Pooled two-sampled Cramer-von Mises test *p* values for 5 Hz evoked first vs second half comparisons (Intra 5 Hz), and comparisons between different 5 Hz stimulus trains (Inter 5 Hz).

**Figure 4.**
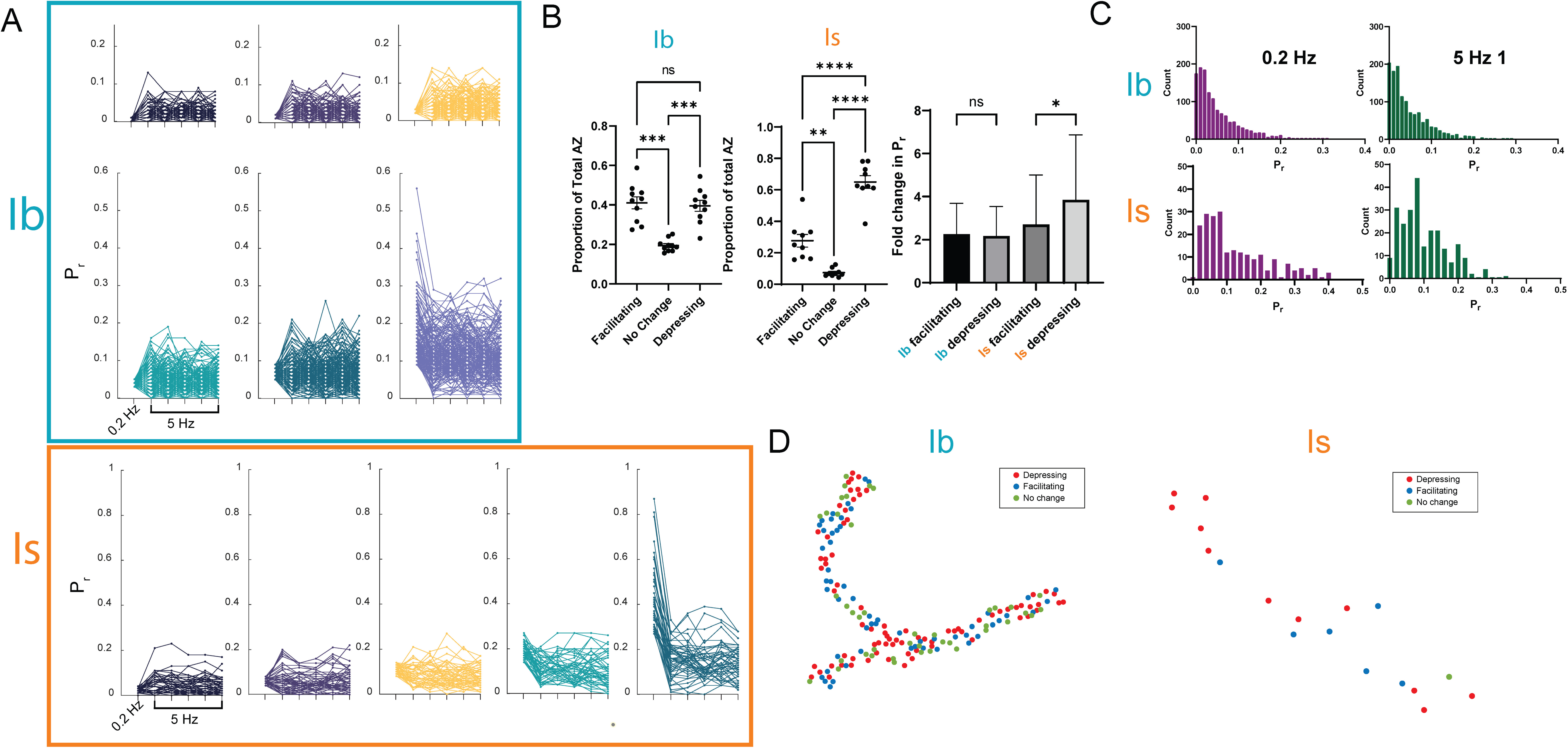
Short-term plasticity varies between active zones with similar basal Pr in both Ib and Is inputs; global vesicular release is constant only for the Ib input. A) *P_r_*change between initial train of 100 stimuli at 0.2 Hz and five subsequent trains of 100 stimuli at 5 Hz across AZs in Ib (top and middle row) and Is (bottom row) axons. AZs binned into six groups by basal *P_r_* at 0.2 Hz (total n = 1304 AZs from N = 10 NMJs; each group contains 217 AZs, except for the highest basal *P_r_* group, which contains 219 AZs). B) Left) Fraction of AZs in each NMJ that facilitate, depress or remain constant upon switch from stimulation at 0.2 Hz to 5 Hz. (Tukey-Kramer post-hoc test: Ib facilitating vs no change, *p =* 0.006; Ib no change vs depressing, *p* = 0.0001; Ib facilitating vs depressing, *p* = 0.9618; Is facilitating vs no change, *p =* 0.007; Is no change vs depressing, *p* < 0.0001; Is facilitating vs depressing, *p* =0.0001). Right) *P_r_*fold change in Ib and Is facilitating and depressing synapses. (Welch’s unpaired t-test, Ib: facilitating vs depressing, *p* = 0.4063; Is: facilitating vs depressing, *p* = 0.001). C) Pooled distributions of *P_r_* for all Ib (light blue) and Is (orange) NMJs (Ib: n = 10 NMJs, 1304 AZs, Is: n = 9 NMJs, 245 AZs) for 100 stimulus train at 0.2 Hz and first 100 stimulus train at 5 Hz. D) Airy-matched coordinates of AZs in a representative Ib and Is NMJ showing spatial distribution of plasticity behavior across the Ib and Is NMJs converging on a common muscle.

**Figure 5.**
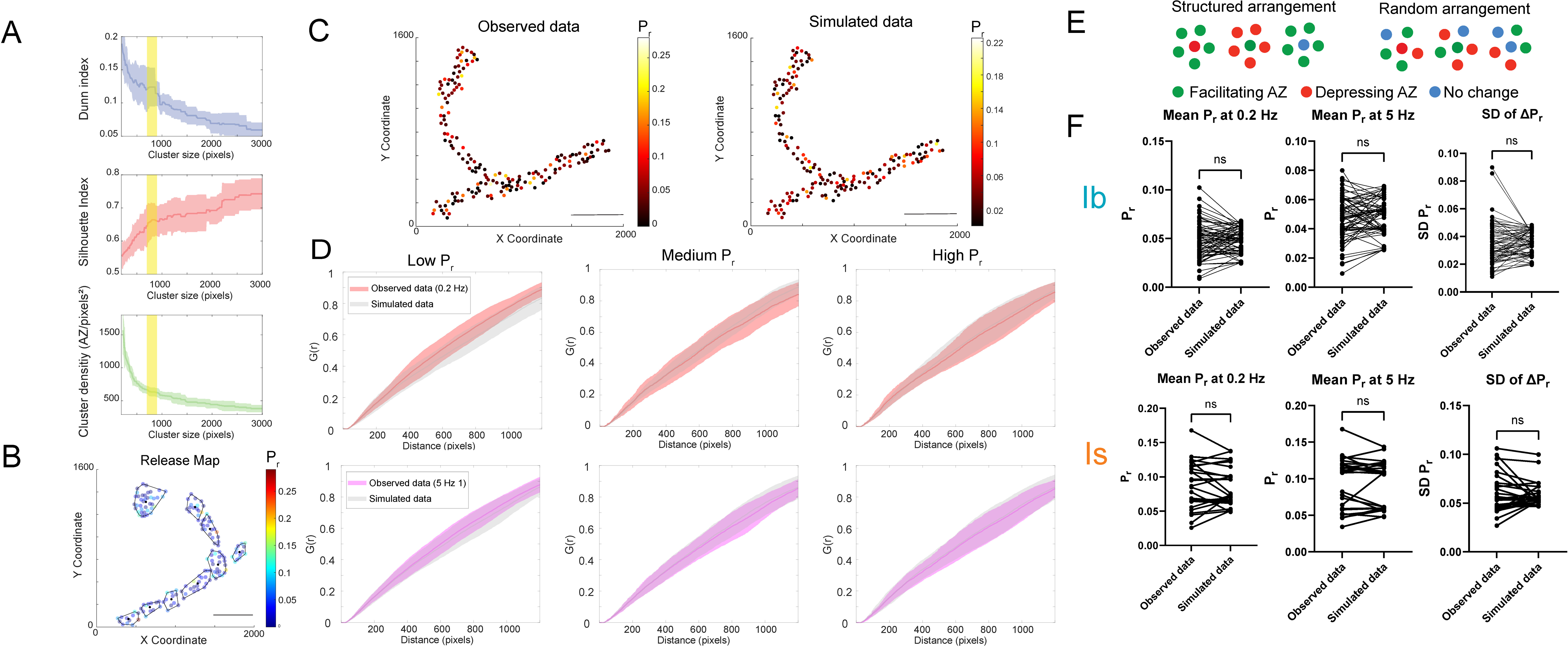
Synapses do not exhibit structured spatial arrangements based on either basal P_r_ or plasticity behavior. A) Cluster analyses detailing three different measures of clustering at various cluster sizes for all ten Ib NMJs. Envelope shows SD across data. The yellow range (700-900 pixels; 23.016 – 29.592 μm) indicates an inflection point in each measure. 1 pixel = 0.03288 μm. B) Example release map with optimal cluster sizes computed from analyses in (A). Each cluster is marked by a boundary. C) Left: Single NMJ example of coordinates of AZ location obtained from Airy-verified molecular imaging. Right) Example of random distribution *P_r_* generated by one iteration of a Monte Carlo simulation. D) Results from distance function (g(d)) analysis conducted on coordinates from three categories of *P_r_*matching three tertiles of total data. Low *P_r_*: 0 ≤ *P_r_* <0.02. Medium *P_r_*: 0.02 ≤ *P_r_* < 0.05, High *P_r_*: *P_r_* ≥ 0.05. Colored lines and envelope represent results computed from observed data, grey lines and envelope represent results computed from simulated data. Results are computed for n = 10 Ib NMJs, simulated results represent n = 10 simulations. Top row: 0.2 Hz stimulation data. Bottom row: first 5 Hz stimulation train data. 1 pixel = 0.03288 μm. E) Diagram showing hypothetical arrangements of AZ based on short-term plasticity behavior. Structured arrangement involves one class of AZ (facilitating, depressing or no change) surrounded by a different class of AZs. Random arrangement involves random plasticity behavior of AZ surrounding a single AZ. F) Plots showing comparison between mean *P_r_* at 0.2 Hz and 5 Hz and SD in the change of *P_r_* of AZ clusters identified via hierarchical clustering for both Ib and Is NMJs. (Two-tailed paired t-tests, Ib: Mean *P_r_*at 0.2 Hz, *p* =0.4328, mean *P_r_* at 5 Hz, *p* =0.3129, SD of change in *P_r_*, *p* = 0.5639, Is: Mean *P_r_* at 0.2 Hz, *p* =0.7054, mean *P_r_* at 5 Hz, *p* =0.7746, SD of change in *P_r_*, *p* =0.7688.

### Constancy of release from a combination of distinct basal release statistics and plasticity diversity

Presynaptic glutamate release sites favor either AP-evoked or spontaneous release,

*i.e.* show a negative correlation between *P_r_* and the frequency of spontaneous release (Peled et al. 2011). In addition, the NMJ exhibits a gradient of synaptic transmission, with weaker proximal boutons and stronger distal boutons, as well as strong branchpoint boutons (Guerrero et al. 2005; Ball et al. 2010). It is not clear how sites with high or low spontaneous or evoked release are distributed spatially the motor nerve terminal. To understand how densities of neurotransmitter release shift across the entire Ib and Is inputs we performed nearest-neighbor analysis of regions of the NMJ, taking each transmission event and computing the probability of another transmission event occurring nearby, within multiple concentric radii. We then adapted a point pattern analysis to the transmission maps. The O-ring statistic (Wiegand & A. Moloney, 2004) was computed and an output of a cumulative distribution function of the k-nearest neighbor distances was analyzed with a Cramer-von-Mise statistical test (**Fig. 3A**). The O-ring statistic computes the expected number of events within some annulus that is a distance *r* from an arbitrary event, providing an estimate of density at different distances. This technique therefore allows us to ask how densities of point patterns change both within boutons and across the entire NMJ.

We initially tested the quality of this statistic by applying it to our data from 0.2 Hz stimulus trains. We compared transmission event coordinates (QuaSOR alone, without structural matching to active zones) in the first 100 and last 100 stimulations. Our premise was that, at this low stimulation frequency, evoked release would be stable and the global spatial statistics should be similar between the first and second halves of the stimulation train. Indeed, we found that overall patterns of release were not significantly different between the first half and the second half of the stimulus train **(Fig. 3B,F**; *p* > α = 0.01). We have previously applied this technique to a comparison of spatial distributions of sites of spontaneous *versus* evoked transmission (Newman et al. 2022). We turned to an analysis of changes in evoked release that take place when animals increase MN firing frequency during locomotion and synapses undergo short-term plasticity. We compared release patterns elicited by trains at 0.2 Hz to ones elicited by trains at 5 Hz, the highest frequency which allowed us to distinguish individual events given the speed of decay of SynapGCaMP6f. We find that for the Ib motor axon, the global patterns are statistically different between the two frequencies **(Fig. 3C).** This suggests that synapses that depress, and thereby reduce their influence on the global statistic, and synapses that facilitate, and thereby increase their influence, are distributed in such a way as to change the global correlation pattern. This effect is absent in the Is motor axon **(Fig. 3G)**, indicating that overall densities of release remain static despite an increase in MN firing frequency. To further understand how these global patterns change, we compared successive 5 Hz trains. We observed little or no statistical difference (Cramer-von-Mise, *p*=α<0.05) within the train (intra-train) or between trains (inter -train) (**Fig. 3E,I**), indicating spatial constancy of global release.

To further characterize the release patterns, we turned from the above global analysis to the analysis of individual synapses. Interestingly, there was no statistically significant difference between the cumulative distribution functions (CDFs) in the 0.2 Hz train and the first 5 Hz train that immediately followed (K-S test, *p* = 0.990) (**Fig. S3 C**). This constancy is striking in view of the changes in global transmission pattern revealed by our Cramer-von-Mise analysis (**Fig. 3)** and the diversity of STP (**Fig. 4)**. With each successive 5 Hz train the number of silent sites increased and comparison of the first 5 Hz train to the fifth showed that the distributions were statistically different (K-S test, *p_1,5_* = 0.007). Together, these results indicate a process by which the overall circuit output is maintained through a constancy of synaptic weights, but where the synaptic weight values have been redistributed among the population of synapses.

### Active zones of similar *P_r_* are randomly distributed in the type Ib MN axon

We sought to understand how *P_r_* and plasticity are spatially distributed across the NMJ. To answer this, we performed a hierarchical clustering analysis using two metrics, the Dunn Index and the silhouette value. The Dunn index is a ratio of the smallest distance between observations not located within the same cluster to the largest intra-cluster distance found within any cluster. The index is used as a metric for evaluating the output of hierarchical clustering, where the result is based on the clustered data itself and does not rely on any external data. The Dunn Index penalizes clusters that have larger intra-cluster variance and smaller inter-cluster variance. In other words, the higher the Dunn index, the better defined the clusters. Typically, the Dunn index favors very small clusters, that is, it assumes the best clusters are individual datapoints. In Ib NMJ boutons, there is a steeper decrease in the Dunn index after ∼900 pixels **(Fig. 5A, top).** We computed the silhouette value for all 10 NMJs. This is a measure of how similar an object is to its own cluster (cohesion) compared to other clusters (separation). The silhouette ranges from −1 to +1, where a high value indicates that the object is well matched to its own cluster and poorly matched to neighboring clusters. If most objects have a high value, then the clustering configuration is appropriate. If many points have a low or negative value, then the clustering configuration may have too many or too few clusters. For Ib NMJs, the silhouette index plateaus after ∼700 pixels **(Fig. 5A, middle).** Thus, both computed clustering metrics indicate a natural cluster forming at the size of an individual bouton and do not detect anything on a smaller scale, indicating absence of clusters at the sub-bouton level **(Fig. 5A, bottom)**.

However, this approach does not answer the question if groups of synapses with similar *P_r_* are grouped together. To address this, we performed a Monte Carlo simulation, using Airy-verified coordinates for AZs **(Fig. 5C)**, and drew random *P_r_* values for each AZ from the exponential distribution fit to the pooled distribution of *P_r_* at 0.2 Hz stimulation (**Fig. 3C**). We wanted to analyze the distance between active zones of similar *P_r_* ranges to determine if AZ location based on *P_r_* followed complete spatial randomness (CSR) or if there was some clustering/dispersion associated with AZs of a similar *P_r_*. In order to test this, we computed nearest neighbor distances based on the g(r)-function (Ripley, 1976), a pairwise correlation function that measures clustering and dispersion of point processes, for all of our simulated iterations as well as our real data.

If active zone location was not dependent on *P_r_* then the results of the g-function of our real data should be constrained within the bounds of our simulated data. For the Ib input, we split our AZs into three groups based on tertiles (groups of three). These ranges were: (Low *P_r_*: 0< *P_r_*≤0.02, Medium *P_r_*: 0.02< *P_r_*≤0.05, and High *P_r_*: 0.05< *P_r_*). We first ran this analysis comparing it to 0.2 Hz data. Remarkably, we found that, in all three groups, at most nearest neighbor distances, the 0.2 Hz g-function was encapsulated by the simulated data (**Fig. 5D, top row**). We repeated this analysis with data from the first 5 Hz stimulation train and found that the computed g-function was also encapsulated by an envelope of simulated g-functions (**Fig. 5D, bottom row).** This analysis was not computed on Is data as there were too few Is AZs to parse AZs into different classes of *P_-r_*. Thus, our Monte Carlo analysis taken together with our hierarchal clustering analysis suggest that AZs of similar *P_r_* are neither dispersed nor clustered, and that within the distance represented by the size of a bouton AZs do not show any structured spatial arrangement based on *P_r_*.

The next step was to determine if groups of active zones within the size of a bouton were organized based on plasticity behavior that changed their *P_r_*in the transition from 0.2 Hz to 5 Hz firing frequency. If spatial location dictates plasticity behavior, then one class of active zone (facilitating, depressing or no change) should be surrounded by a different class of active zone (**Fig. 5E**). If the spatial arrangement were random, then both the mean *P_r_*and standard deviation of *P_r_* within clusters should be statistically different from randomly simulated data. To test this, we performed hierarchical clustering based on spatial distance between active zones. We then computed the mean *P_r_* within clusters for both 0.2 Hz and 5 Hz. A Monte Carlo simulation was then performed by shuffling *P_r_* across the entire Ib or Is input and then computing the mean *P_r_* within those same clusters. For both Ib and Is inputs, mean *P_r_*of clusters at 0.2 Hz and 5 Hz was not statistically different from simulated data (Two-tailed paired t-tests, Ib: Mean *P_r_* at 0.2 Hz, *p* =0.4328, mean *P_r_* at 5 Hz, *p* =0.3129, SD of change in *P_r_*, *p* =0.5639, Is: Mean *P_r_* at

0.2 Hz, *p* =0.7054, mean *P_r_* at 5 Hz, *p* =0.7746, SD of change in *P_r_*, *p* =0.7688) (**Fig. 5F**). These results suggest that, while global densities of release may exist from a proximal to distal gradient across an entire axon (Ball et al. 2015), local clusters of synapses within a bouton do not exhibit a specific spatial arrangement based on either basal *P_r_* or plasticity behavior.

### The Drosophila metabotropic receptor regulates transmitter release in an input-specific manner

Our observations indicate that the Ib MN input to muscle is highly plastic, and that the Is MN input is more static. Previous studies have proposed that glutamate in the synaptic cleft could serve as a homeostat to regulate release as to maintain it at an optimal level (Daniels et al., 2004; Li et al., 2018). Such a system needs to sense glutamate concentration, perhaps by presynaptic auto-receptor GPCRs. In *Drosophila*, the sole metabotropic glutamate receptor (DmGluRA) has been proposed to be expressed in the presynaptic MN and not muscle (Budnik et al. 2004).

To determine if DmGluRA could contribute to the diversification of release properties, we employed the Gal4/UAS system to knock down DmGluRA expression in type I MNs using an RNAi. We QuaSOR mapped transmission events evoked by 1 Hz stimulation. Comparison of the UAS-DmGluRA-RNAi knockdown to an attP2 insertion site control (UAS-attP2) showed that quantal density in Ib and Is did not differ between the knockdown and control at either 0.2 Hz and 5 Hz MN stimulation (**Fig. 6A**) (2-tailed repeated measures ANOVA test with Greenhouse-Geiser correction, *p*-values: 0.2 Hz: attP2 vs DmGluRA-RNAi Ib, *p*=0.7598, attP2 vs DmGluRA-RNAi Is, *p*=0.7927, 5 Hz: attP2 vs DmGluRA-RNAi Ib, *p*=0.8490, attP2 vs DmGluRA-RNAi Is, *p*=0.8138). The lack of effect of receptor knockdown on release at these stimulation frequencies is consistent with what was reported earlier based on electrophysiological analysis of stimulation at up to 20 Hz (Budnik et al. 2004) and suggests that DmGluRA may function at higher frequencies of firing (>40 Hz), such as those that occur during locomotion (Kadas et al., 2017). Since we could not perform QuaSOR in contracting muscle, to assess the effect of DmGluRA activation on release, we turned to application of the mGlur2/3 agonist LY354740. LY354740 inhibited global synaptic release at type Ib synapses but not type Is synapses **(Fig 6B,C)** (Kolmogorov-Smirnov test, *p*-values: attP2 Ib, *p* <0.0001, attP2 Is, *p* = 0.225, DmGluRA-RNAi Ib, *p =* 0.003, DmGluRA-RNAi Is, *p* = 0.221). The lack of effect at Is synapses may be explained by the lower level of Is expression of DmGluRA reported in RNAseq analysis (Jetti et al., 2023). The LY354740 inhibition of release in type Ib synapses was most pronounced at ones with high *P_r_* **(Fig. 6D)**. These are the synapses that we found, above, to depress at moderately elevated firing frequency **(Fig. 4A)**. These findings suggest that the synapses with the highest basal synaptic weights dominate transmission at low action potential firing frequencies and play an increasingly smaller role as firing frequency increases, first because of an intrinsic tendency toward short-term depression and second, at the high firing frequencies that mediate locomotion, because of feedback inhibition by released glutamate.

**Figure 6:**
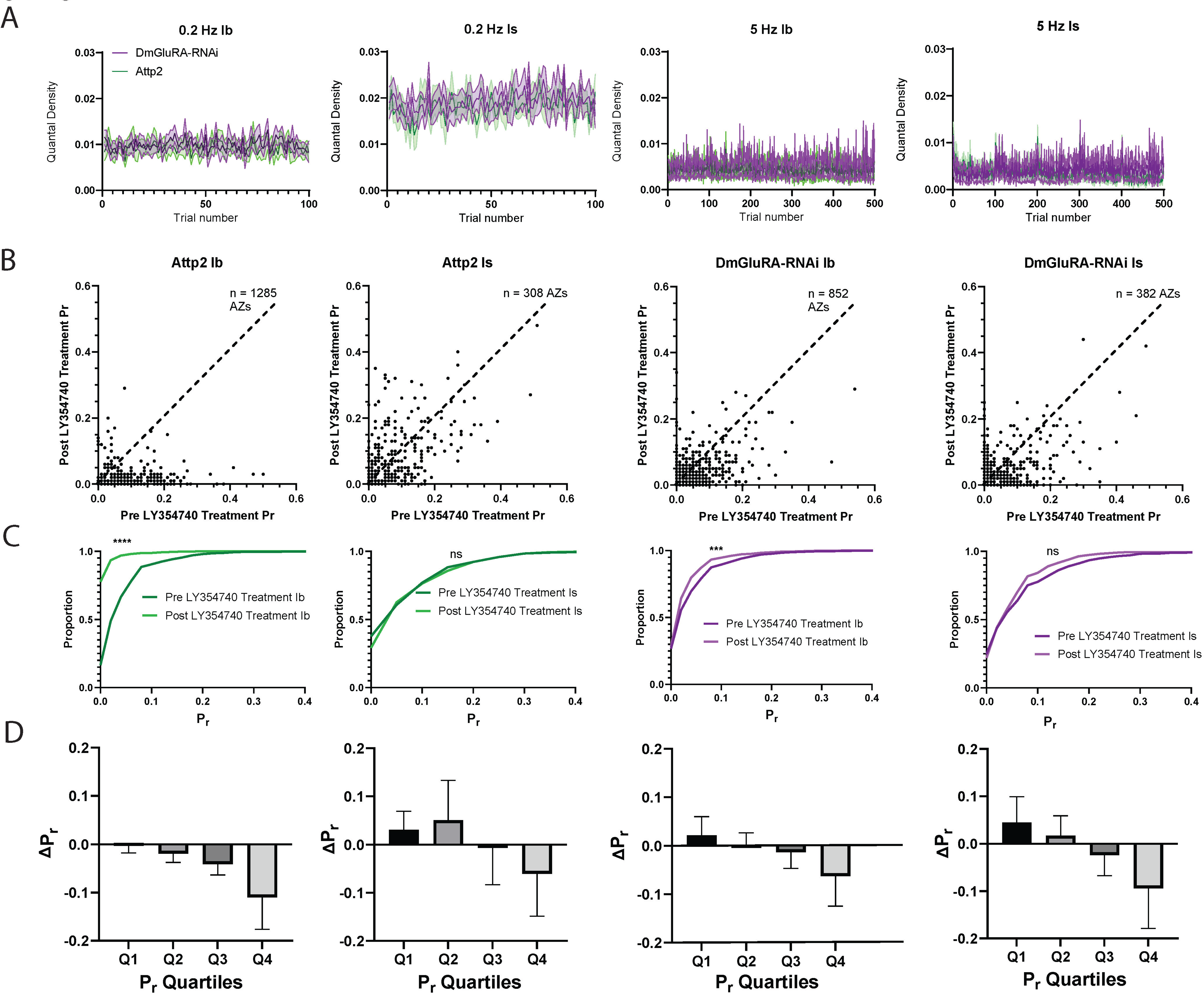
DmGluRA selectively inhibits release from Ib but not Is synapses. A) Quantal density measurements for DmGluRA-RNAi and Attp2 control animals for both Ib and Is inputs. Top row: quantal density measured during basal (0.2 Hz) evoked release. Bottom row: quantal density measured during high frequency (5 Hz) evoked release. Upper and lower bounds indicated by dashed lines, shaded shows range (n = 5 animals for each genotype). B) Plots showing individual AZ *P_r_* before and after application of DmGluRA agonist, LY35470 at 50 1M. Pre-LY3570 *P_r_*on the x-axis and post-LY35470 *P_r_* on the y-axis. C) Cumulative distribution plots of *P_r_* before and after application of LY354740 (Kolmogorov-Smirnov test, *p*-values: Attp2 Ib, *p* <0.0001, Attp2 Is, *p* = 0.225, DmGluRA-RNAi Ib, *p =* 0.003, DmGluRA-RNAi Is, *p* = 0.221). D) Plots showing raw changes in *P_r_* following application of LY354740 for different quartiles based on starting *P_r_*.

## Discussion

Both in vertebrates and invertebrates, synapses made by an individual neuron can vary between high and very low *P_r_* (del Castillo & Katz, 1954; Dobrunz & Stevens, 1997; Hessler et al., 1993; Katz & Miledi, 1967; Mendonça et al., 2022; Newman et al., 2022)). Such differences in synaptic weight can arise developmentally or from activity-dependent plasticity (Atwood & Karunanithi, 2002; Han & Stevens, 2009; Harris & Littleton, 2015; Medeiros et al., 2024) While there is a general tendency, during AP firing activity at elevated frequency, for high *P_r_* synapses to depress, due to depletion of releasable vesicles, and for low *P_r_* synapses to facilitate (Peled & Isacoff, 2011), it has not been clear if there are additional mechanisms that diversify synaptic plasticity. We find that synapses formed by an individual *Drosophila* larval MN vary greatly in short-term plasticity, even when they start off at similar basal *P_r_* and even though they all synapse onto a common postsynaptic muscle fiber.

Our analysis shows that synapses are not clustered according to either basal *P_r_* or the direction (facilitation *versus* depression) or degree of their short-term plasticity; nor is there a tendency for strong and weak or facilitating and depressing synapses to be located near one another in a way that would suggest competition. This suggests that the presynaptic plasticity determinants are localized to—and operate separately at—each active zone and that they are distributed randomly in the presynaptic terminal.

However, despite the apparently random distribution of synapses with distinct basal strength and plasticity, overall transmission strength among the population of synapses of a type Ib MN remains constant when the neuron’s firing frequency changes and diverse plasticity changes occur across the diverse synapses. This constancy in synaptic weight distribution emerges from a balance between the number of synapses that facilitate and the number that depress, their degree of plastic change and the distribution of their initial basal *P_r_*. Type Is MN synapses also have a limited degree of balancing and are dominated by depression and a narrowing of *P_r_* distribution at elevated firing frequency.

Stochasticity is an integral part of the nervous system (Heams, 2014; Ilan, 2020; W.-H. Zhang et al., 2023), underlying and often enhancing function at multiple biological scales. The biological reason for such spatial randomness has been explored in theoretical studies (Arleo et al., 2010; Manwani & Koch, 1997, 2001; C. Zhang & Peskin, 2015). Some reasons include inherent stochasticity arising from development (Atwood & Karunanithi, 2002; Han & Stevens, 2009) and enhancement of transmission that allows a neuron to better discriminate between varying firing frequencies. Other studies have highlighted the importance of randomness in maintaining information transfer across the synapse, modeling this process through the use of optimal linear filter theory, information theoretic measures, and continuous-time Markov chains (Goldman, 2004; Reich & Rosenbaum, 2013; C. Zhang & Peskin, 2015). For example, *in silico* simulations show randomness allows a neuron to prioritize information flow through higher-frequency presynaptic spike trains, and may also allow for optimum energy conservation (Malkin et al., 2024; Schug et al., 2021). Understanding how a postsynapse integrates information from the presynapse when: 1) vesicle release is stochastic and 2) presynaptic parameters such as spatial location, firing frequency of the presynaptic input, and molecular composition of the active zone change over time therefore requires the study of a system where synaptic release behavior can be measured simultaneously and under multiple presynaptic conditions.

The notion that synapses are able to balance synaptic strength among groups of synapses is consistent with the theory of homeostatic synaptic scaling, wherein a neural network adjusts parallel synaptic pathways and excitability of a postsynaptic neuron to compensate for individual changes to synaptic strength in such a way that the overall firing rate/output of that network is maintained even when some synapses are strengthened or weakened (Harris-Warrick & Marder, 1991; Marder & Goaillard, 2006; G. Turrigiano, 2011; G. G. Turrigiano, 2008b; G. G. Turrigiano et al., 1998). One mechanism by which specific synapses restore potency of transmission following changes that affect one side of the synapse is presynaptic homeostatic plasticity (PHP), in which transmitter release is adjusted to compensate for changes in postsynaptic sensitivity to transmitter (Davis & Goodman, 1998; Frank, Davis, 2004, Davis & Müller, 2015). However, homeostatic synaptic scaling occurs over hours and PHP over minutes, whereas the synaptic re-balancing that we observe here adjusts within the time it takes for synapses to depress or facilitate to new steady-state levels, i.e. within seconds. Thus, the nervous system appears to have several mechanisms that operate over different time scales to maintain balance and constancy of circuit output across changing conditions.

Synaptic re-balancing is accompanied by marked changes in the spatial distributions of the AZs that release the most neurotransmitter. What mechanisms could underlie this phenomenon? A candidate mechanism that could alter the release probabilities of dozens of synapses on a large scale is the presynaptic auto-receptor pathway (Davis & Müller, 2015; Langer, 2008). Typically, autoreceptors are located perisynaptically on the presynapse and are G-protein coupled receptors (GPCRs) that bind the neurotransmitter released from the presynapse (Langer, 2008). At the Drosophila NMJ, there is a single presynaptic metabotropic glutamate receptor, the autoinhibitory DmGluRA, whose activation suppresses glutamate release (Bogdanik et al., 2004). Negative feedback through DmGluRA could dynamically regulate *P_r_*and do so in a differential manner that depends on level of expression of DmGluRA itself or components of its downstream signaling cascade, which include CaMKII and PI3K (Howlett et al., 2008). Previous studies (Daniels et al., 2004; Li et al., 2018) have proposed extracellular glutamate concentration as a possible homeostatic set point, and the DmGluRA pathway might represent one such pathway to homeostatically regulate release. Here, we show that, consistent with previous findings (Bogdanik et al., 2004), knockdown of this receptor does not change transmission at 0.2 Hz or 5 Hz stimulation. However, a DmGluRA agonist produces presynaptic inhibition, suggesting that at higher frequencies of activity, as occur during locomotion (Kadas et al., 2017), DmGluRA activation may be sufficient to provide autoinhibition and limit release. Interestingly, agonist inhibition only occurs in the Ib MN, not the Is MN, and with the most pronounced effect in highest *P_r_*active zones. This indicates that, as presynaptic firing frequency changes so too does the distribution of synaptic weights that dominate at a specific frequency. Thus, our findings suggest that synapses with high basal synaptic weights dominate transmission at low action potential firing frequencies and play an increasingly smaller role as firing frequency increases, first because of an intrinsic tendency toward short-term depression and second, as firing frequency rises further during locomotion, because of feedback inhibition by released glutamate. Over the low to mid-range in firing frequency range that precedes substantial contraction, where we were therefore able to quantify synaptic weight distribution across all of the synapses formed by the type Ib MN, the reduced impact of high *P_r_* active zones is replaced by increased contribution from low *P_r_* active zones, providing remarkable constancy. In contrast, the Is MN population of AZs depresses more than it facilitates and thus its drive for locomotion reduces in relation to that of the type Ib input. This suggests that, as locomotory drive increases, tonic ramp-burst transmission of the type Ib MN, becomes more dominant than the phasic abrupt burst transmission of the type Is MN.

Homeostatic adjustments in the nervous system in response to developmental changes, activity-dependent plasticity changes and changing environmental conditions have been shown to regulate both neuronal firing and synaptic transmission over a time-course of minutes to hours. The adjustments can have multiple solutions, *i.e.* accommodate to individual neuronal variability to obtain global constancy of circuit output (Marder et al. 2023). Here we find that the variability extends to individual synapses, the constancy to the summed output of the synaptic population and the rate of adjustment to occur over seconds. This constancy in output across changes in presynaptic firing could enable reliable behavior in the face of different environmental challenges.

## Methods

### Fly Genetics

Flies were raised on standard corn meal and molasses media at 25°C. Wandering third instar larvae were used in all experiments. For UAS-DmGluRA-RNAi animals, third instar larvae were screened using an Axio Zoom.V16 microscope (Carl Zeiss, Oberkochen, Germany) using balancers with larval markers including CyO^GFP^ (3xP3-EGFP variant) and TM6B. Only actively crawling larvae were used for experiments. Wild-type heterozygous SynapGCaMP6f was used for analysis (w^118^;+;+/SynapGCaMP).

### Functional Imaging

Optical quantal imaging was performed similarly to our previous report. Briefly, third instar larvae were dissected on PDMS (Sylgard 184, Dow Corning, Auburn, MI) pads in ice-cold HL3 solution containing, in mM: 70 NaCl, 5 KCl, 0.45 CaCl_2_ · 2H_2_O, 20 MgCl_2_ · 6H_2_O, 10 NaHCO_3_, 5 trehalose, 115 sucrose, 5 HEPES, and with pH adjusted to 7.2. Following removal of the brain, larval fillets were washed and imaged in room temperature HL3 containing 1.5 mM Ca^2+^ and 25 mM Mg^2+^.

Fluorescence images were acquired at room temperature with a Vivo Spinning Disk Confocal microscope (3i Intelligent Imaging Innovations, Denver, CO), using a 63 × 1.0NA water immersion objective (Zeiss) mounted on an upright Examiner Zeiss Microscope, 1.2X optical adapter, LaserStack 488 nm (50 mW) laser, CSU-X1 A1 spinning disk (Yokogawa Tokyo, Japan), standard GFP filter, and EMCCD camera (Photometrics Evolve512, Tucson, AZ). All live SynapGCaMP6f imaging recordings were done on ventral longitudinal abdominal muscle 4 at segments A3-A5 of third instar larvae. All imaging was performed using 50 ms exposures (20 fps) of the full camera sensor (512 × 512 px).

Nerve stimulation was performed with a suction electrode attached to a Stimulus Isolation Unit (SIU, ISO-Flex, A.M.P.I Jerusalem, Israel) with 100 μs stimulus duration. Stimulation intensity was adjusted to recruit both Ib and Is axons, as verified during the imaging. Nerve stimulation and imaging were synchronized using custom-written MATLAB scripts (MATLAB Version 2015b, MathWorks, Inc., Natick, MA) in order to control the SIU and trigger imaging episodes within SlideBook (v6.0.16, 3i Intelligent Imaging Innovations).

To gather evoked transmission events at each NMJ, two separate quantal imaging experimental protocols were utilized. In the sequential protocol, we collected AP-evoked responses during short, single-stimulus episodes. Here each stimulus was collected during a series of 10 images (50 ms exposures). Each episode had 3–4 baseline frames prior to nerve stimulation. We collected either 100 or 200 single-stimulus episode trials at 0.2 Hz. This was followed immediately by spontaneous event collection, imaging continuously at 20 FPS (50 ms exposures in streaming capture mode) for 2 minutes total separated into four, 30 s movies. These brief pauses between movies allowed for the manual correction of any drift in the NMJ. In the second interleaved activity imaging protocol, both spontaneous and evoked events were collected during the same image acquisition protocol. This was done by imaging continuously at 20 FPS (50 ms exposures in streaming capture mode) for 30 s while stimulating the nerve once per 200 ms during each 30 s movie (5 Hz). Data was also acquired this way while stimulating the nerve once per second for 100s (1 Hz). Minor adjustments were made between movies to account for drift and muscle contraction, with a maximum of 10 s between movies. To ensure comparability between experiments, recordings were done on only one NMJ per larva (i.e. the number of NMJs = the number of larvae) in recordings that were performed within 30 min of the beginning of the dissection, thereby ensuring health and similar conditions. For experiments with the DmGluRA agonist, LY354740 (Tocris Bioscience, No.3246, Minneapolis, MN) was resuspended in a molar equivalent of NaOH at 50 mM. After an initial 1 Hz recording, the agonist was added to a still bath by pipetting an equal volume of a 2x working stock solution, diluted in the appropriate recording HL3 saline. The large volume was used to ensure rapid mixing of the solution without continuous perfusion. A second 1 Hz recording was taken 15 minutes after application of LY354740. This time period allowed for equilibration and penetration of the drug into live tissue.

### Immunohistochemistry

Larvae were fixed in room temperature Bouin’s fixative (Ricca Chemical Company, Arlington, TX) for 5 min, permeabilized in PBS with 0.1% Triton X100 (PBT) and blocked in PBS with 0.1% Triton X100, 5% normal goat serum, and 0.02% sodium azide (PBN). All antibody incubations were performed in PBN and all washes were performed in PBT. Mouse anti-Brp (nc82; Developmental Studies Hybridoma Bank, Iowa City, IA) was used at 1:100 for Airyscan imaging. Chicken anti-GFP (Thermo Fisher A10262; Thermo Fisher Scientific Waltham, MA) or anti-GFP FITC conjugated (Rockland 600-102-215) were used at 1:1000 to label SynapGCaMP6f in fixed samples. Alexa Fluor 647 (Jackson 123-605-021) and Cy3-conjugated goat anti-Hrp (Jackson 123-165-021; Jackson ImmunoResearch Laboratories, West Grove, PA) antibodies were used at 1:250. Alexa Fluor 405 goat anti-mouse (Thermo Fisher A31553) or Alexa Fluor Plus 555 goat anti-mouse (Thermo Fisher A32727), Alexa Fluor 488 goat anti-chicken (Thermo Fisher A11039) were used at 1:1000.

### Airyscan imaging and analysis

Following antibody incubations and washes, larval fillets were mounted in Vectashield (H-1000; Vector Laboratories, Burlingame, CA), Vectashield HardSet (H-1400), or Vectashield Vibrance (H-1700). Confocal Airyscan imaging was performed on either a Zeiss LSM 880 or Zeiss LSM 980 microscope. All samples were imaged with a ×63 oil immersion objective (NA 1.4, DIC; Zeiss) using Zen software (Zeiss Zen Blue 3.5, 3.7). All imaging data were collected using identical imaging and processing parameters for a given experiment set. Unless otherwise noted all confocal and Airyscan images are displayed as Gaussian filtered maximum intensity projections that were generated using custom-written MATLAB routines.

Airyscan imaging data were acquired using a tiling strategy, whereby smaller volumes of each NMJ were acquired sequentially and stitched together. Brp reconstructions for matching to QuaSOR data were acquired on the LSM 880 system. Briefly, each imaging volume was acquired with an additional magnification of 12x with a 5 AU pinhole, 2 µs pixel dwell times, line averaging of 2, an *x-y* dimension of 1024 by 1024 px (processed to 1000 by 1000 px) at 11 nm/px, and axial *z* spacing of 159 nm. Each of the three-channel volumes (anti-Brp/Alexa 405, anti-GFP/Alexa 488, and anti-Hrp/Cy3) were scanned sequentially. Alexa 405 was excited with a 405-nm laser and data were acquired with a BP420-480-LP605 filter. Alexa 488 was excited with a 488-nm laser and data were acquired with a BP495-550 + LP570 filter. Cy3 was excited with a 561 laser and data were acquired with a BP495-550 + LP570 filter. Airyscan processing of all channels and *z* slices was performed in Zen (Zeiss Zen Black v2.3 SP1) using super-resolution settings. Additional Brp intensity quantification data were acquired on an LSM 980 system with 4x magnification, a 5 AU pinhole, 0.77 µs pixel dwell times, an x-y dimension of 1024×1024 px (processed to 1000×1000) at 32 nm/px and axial z spacing of 130 nm. Here anti-GFP conjugated to FITC was excited with 488 nm and SP550 filter, anti-Brp/Alexa Fluor Plus 555 was excited with 561 nm and SP615 filter, and anti-HRP Alexa Fluor 647 was excited with 639 nm and LP570 filter, and images were line scanned bidirectionally with no line averaging. Airyscan processing was conducted using Zeiss Joint Deconvolution (jDCV) software. We then sequentially stitched each 3D volume together in Fiji (NIH ImageJ Version 2.0.0-rc-43/1.52n) using pairwise stitching with linear blending. Alignments were performed to maximum intensity pixels of Brp puncta in overlapping volumes. Regions outside of the tile borders are always indicated by gray coloring in the corresponding images. AZ locations in stitched Airyscan datasets were calculated by masking the volumetric Brp data. Sites were initially identified using a local 3D Brp intensity maximum with a minimum distance of 150 nm from neighboring maxima. AZ identifications were manually validated and corrected when neighboring sites were misidentified. AZ-specific Brp voxel intensities were calculated by identifying 3D-connected voxels to each AZ maxima for isolated AZs.

### Quantal super-resolution optical reconstruction (QuaSOR)

All quantal super-resolution optical reconstruction (QuaSOR) analysis was performed as described in our previous work (Newman et al 2022). Briefly, raw images were converted, bleach corrected, and registered for movement in the x-, y- and z-axis. Once images were registered event detection via pixel correlation and thresholding was performed. Images were considered to be events if they were above a certain dF/F threshold, usually 0.04/0.05dF. These events were then compared to overall response fields to eliminate false positives. QuaSOR was then performed on extracted events. This involved matching groups of pixels to a mixture-of-models series of Gaussians. Once an appropriate match was found the event was refined and event coordinates were produced. Structure-function matching analysis is detailed in our previous work (Newman et al., 2022).

### Computation and statistics Clustering analysis

***O-ring statistic.*** To assess the relative co-localization between different sets of QuaSOR coordinates, we adapted a technique for the analysis of ecological point pattern processes (Wiegand & A. Moloney, 2004) termed the O-ring statistic. Firstly, coordinates of all Ca^2+^ transmission events under a particular stimulation condition are obtained and projected onto an xy-plane. Then a k-NN metric is computed for all events. This k-NN distance is then used as input as part of a Ripley’s-K distance function K(r) (Ripley, 1976), which computes the expected number of points in a circle of radius r centered at an arbitrary point. An intensity *I* of all points is then computed and the alternative pair correlation function g(r) is then used. This function g(r) replaces the circles associated with Ripley’s-K function with rings, and ultimately give the expected number of points at some distance r from an arbitrary point divided by the intensity I of the Ca^2+^ transmission events. This is termed the O-ring statistic. The O-ring statistic is closely related to the K-function. The K-function examines how points in space are correlated and as its output computes the expected number of events within a radius r from an arbitrary event. The O-ring statistic builds upon this but computes the expected number of events within some annulus that is a distance r from an arbitrary event, providing more accurate estimates of density at different distances than the K-function. In this scenario, the output of the O-ring statistic would be the average number of transmission events some distance r away in an annulus from an arbitrary transmission event.

This O-ring statistic used distances between points to determine the neighborhood density of transmission events. A probability density function (PDF) detailing the neighborhood density of points within an annulus distance r away from an arbitrary point is generated. An empirical cumulative distribution function is then generated from this PDF, with distances (x-axis) scaled from 0 to 1. These scaled distances are termed “transformed distances”. 0 represents the origin of a point at the center of an analysis whereas 1 represents the maximum possible distance within the bounds of an NMJ that a point can be away from the center point. From these eCDFs a comparison is made using the two-sample two-sided Cramér–von Mises test to examine differences in distances between the comparison groups. Statistically significant differences in distances between conditions likely indicated a distinct underlying point process for the coordinate sets. Algorithm was written using R.

#### Dunn Index

The Dunn Index is computed as follows for given *m* clusters:

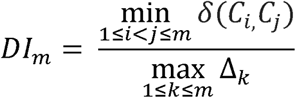

Where δ(*C_i,_C_i_* ) is the distance computed between any two points out of *k* points.

#### Silhouette value

The silhouette value is defined as follows through a silhouette coefficient (SC):

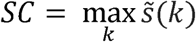

Where *s̃*(*k*) represents the mean *s(i)* over all data of the entire dataset for a specific number of clusters *k*.

#### G-function

The G-function is computed as follows:

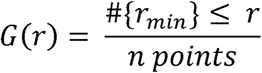

The output is thus a cumulative distribution function that examines the nearest neighbor distances at some minimum distance r_min_ away.

### Monte Carlo Simulation

All Monte Carlo simulations were performed on QuaSOR-Airy matched coordinates using custom scripts written in MATLAB (MATLAB Version 2021b, MathWorks, Inc., Natick, MA). Underlying probability distributions for *P_r_* were determined via fits of several different probability distributions to histograms of *P_r_* (not shown) and used maximum-likelihood estimation (MLE) and Aike information criterion (AIC) to test the goodness of fit. The exponential distribution was found to have the maximal log-likelihood (3823) and minimal AIC score (-5764) when both tests were conducted on the 0.2 Hz *P_r_*distributions. Probabilities of release were thus simulated using underlying exponential distributions computed via Maximum Likelihood Estimation on individual distributions of 0.2 Hz *P_r_*. The formula for the exponential probability density functions was as follows:

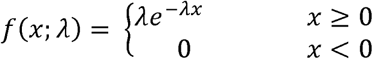

Average λ for all fit exponential distributions was computed, and a sample distribution was established. AZ coordinates were then indexed randomly, and for each x-y coordinate pair a sample *P_r_* was drawn from the sample distribution and assigned. This process was iterated for at least 100 times for each comparison and for each stimulation condition (0.2 Hz, 5 Hz). For each iteration a distance function g(r) was computed (Ripley, 1976), and the resulting function was plotted and compared to distance functions generated using ground-truth data.

## Code availability

Code used for all spatial and statistical analyses will be available in a Github repository upon publication. QuaSOR custom code is available in the GitHub repository under the filename https://github.com/newmanza/Newman_QuaSOR_2021; https://doi.org/10.5281/zenodo.5711302.

## Data availability

All imaging, statistical and analytical data will be available in a Github repository upon publication.

## Acknowledgements

We thank Amy Winans, Victoria Shih-Wei Chou, Adam Hoagland and Robin Ball for their helpful discussions. We thank the rest of the Isacoff lab for their helpful comments. We thank Marla Feller for her valuable suggestions and discussion. We thank Feather Ives and Holly Aaron of the Berkeley Microscopy Imaging Center (MIC) for installing and maintaining the experimental microscopes. The study was supported by the National Institutes of Health Grant No. R01NS107506 (E.Y.I.) and a UCLA National Institute of Health Institutional Research and Academic Development Award 5K12GM106996 ( K.A.). E.Y.I. is a Weill Neurohub Investigator.

## Contributions

K.A., R.S., R.L., and Z.L.N performed optical quantal imaging and QuaSOR analysis. D.B wrote code for and performed further QuaSOR analysis. P.M. wrote code for and performed statistical analyses. R.S., R.L. and K.A. performed antibody staining, Airyscan imaging and analysis. Z.L.N., K.A. and E.Y.I designed the functional imaging experiments. K.A. and E.Y.I. wrote the paper with input from all of the authors.

