## Supplemental Figure Legends for "Synapse-to-synapse plasticity variability balanced to generate input-wide constancy of transmitter release"

### Supplemental Figures Legends

### Figure S1, related to Figure 1. QuaSOR super-resolution mapping of sites of evoked transmission.

A) Single AP-evoked ∆F response frame. Gray regions indicate areas of the muscle that were not associated with either Ib or Is NMJ and therefore not analyzed. ∆F range: 0 – 13686

AU.

B) ∆F/F for entire NMJ after bleach correction and amplitude thresholding.

C) ∆F/F corresponding to boxed region in (B).

D) Same events as in (C), showing events with isolated ∆F/F after cleanup (B) showing isolated ∆F/F (only pixels associated with event).

E) QuaSOR-matched Gaussians applied to 7 events isolated in (D). Maxima for 2D Gaussians are indicated by black dots at the center of the Gaussian (dark red). ∆F/F for C – E: 0 – 0.97. Scale bars: 10 µm (A, B), 2 µm (C – E).

### Figure S2, related to Figure 2. Transmission dynamics of identified synapses.

A) Partial dataset of transmission dynamics at structurally identified AZ locations for Ib NMJs 3 and 8. Red in NMJ#3 and NMJ#8: AZs labeled according to index from #1-32 (Top, NMJ#3) and from #138-160 (Bottom, NMJ#8).

B) Zoomed in subset of AZs indexed in (A). Top: Each AZ is assigned an index. Bottom: Raster plot tracking AP-evoked quantal release behavior of each AZ. Stimulus # during experiment is on the x-axis, AZ ID is on the y-axis. Each dot corresponds to an event assigned to an index AZ after QuaSOR-Airy structure-function matching.

C) Full raster plots for all identified AZs in Ib 10 NMJs. Blue: 0.2 Hz, Red: 5 x 5 Hz.

### Figure S3, related to Figure 4. P_r_ constancy across stimulation frequency.

A) Each subpanel represents a single Ib NMJ and its distributions of P_r_ collected through multiple stimulation modalities (0.2 Hz, five 5 Hz trains). In purple are the distributions for the 0.2 Hz evoked modality, and green distributions are for the 5 Hz trains. X-axes are normalized across the NMJs but y-axes are proportional to within NMJ AZ count.

B) Pooled distributions of P_r_ for all NMJs (n = 10 NMJs, 1304 AZs). Purple is 0.2 Hz modality and green are the 5 Hz modalities. Black lines are exponential fits to each distribution.

C) Cumulative distribution function (CDF) comparing pooled P_r_ between 0.2 Hz modality and first 5 Hz modality. D) CDF comparing all five 5 modalities.
