## Supplementary figures and images for "Synapse-to-synapse plasticity variability balanced to generate input-wide constancy of transmitter release"

### Supplemental Figure 1

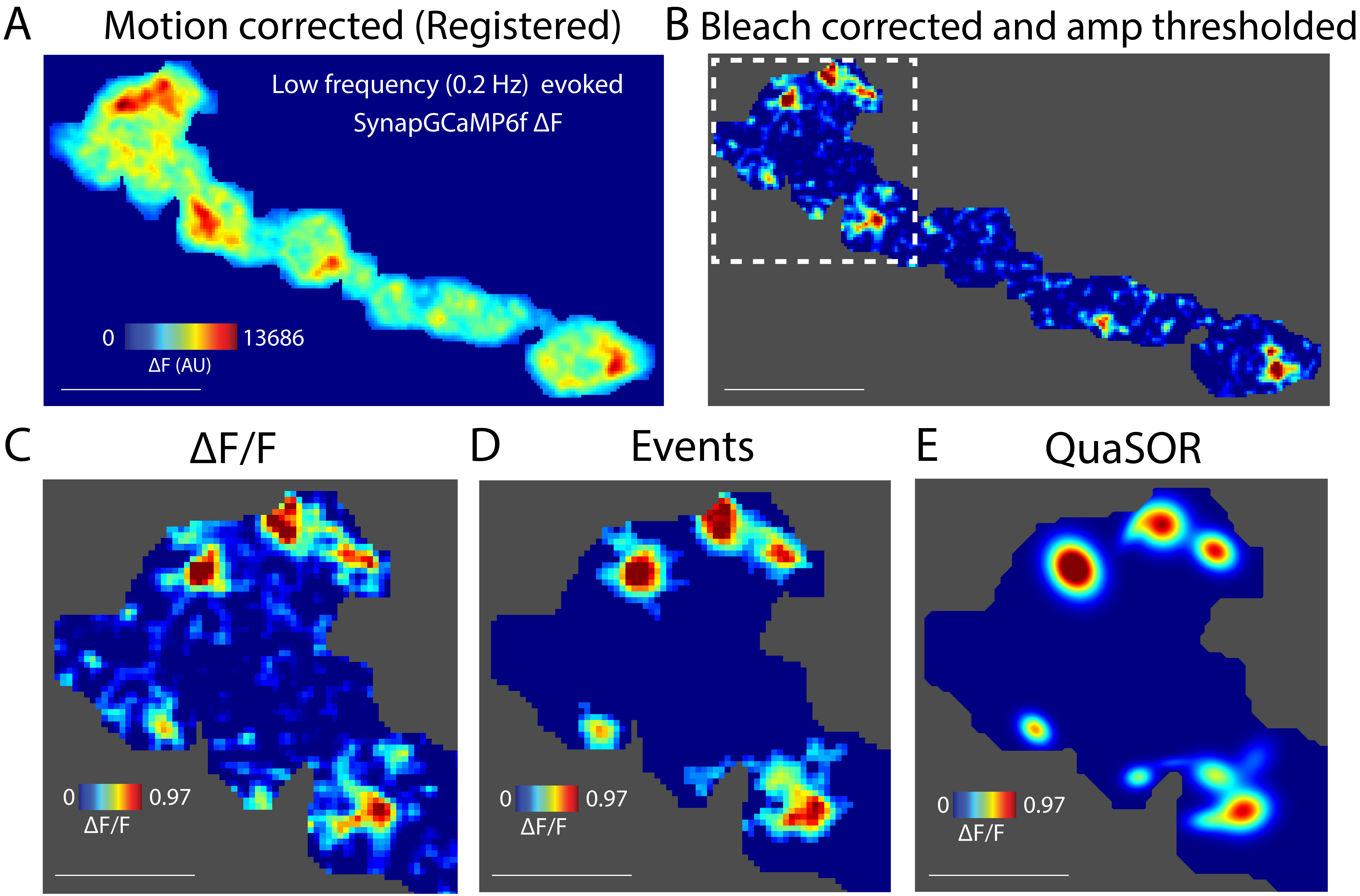

### Supplemental Figure 2

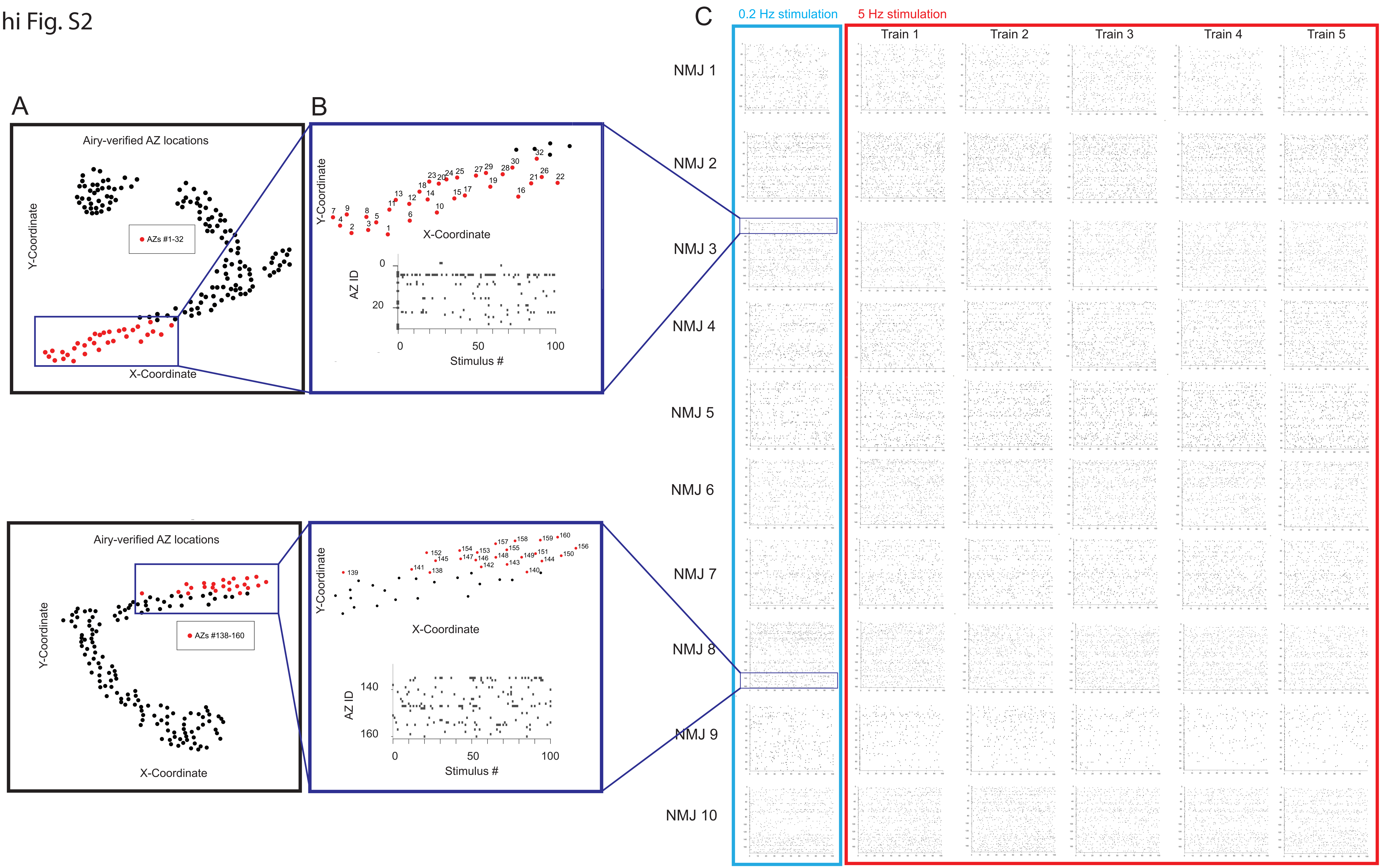

### Supplemental Figure 3

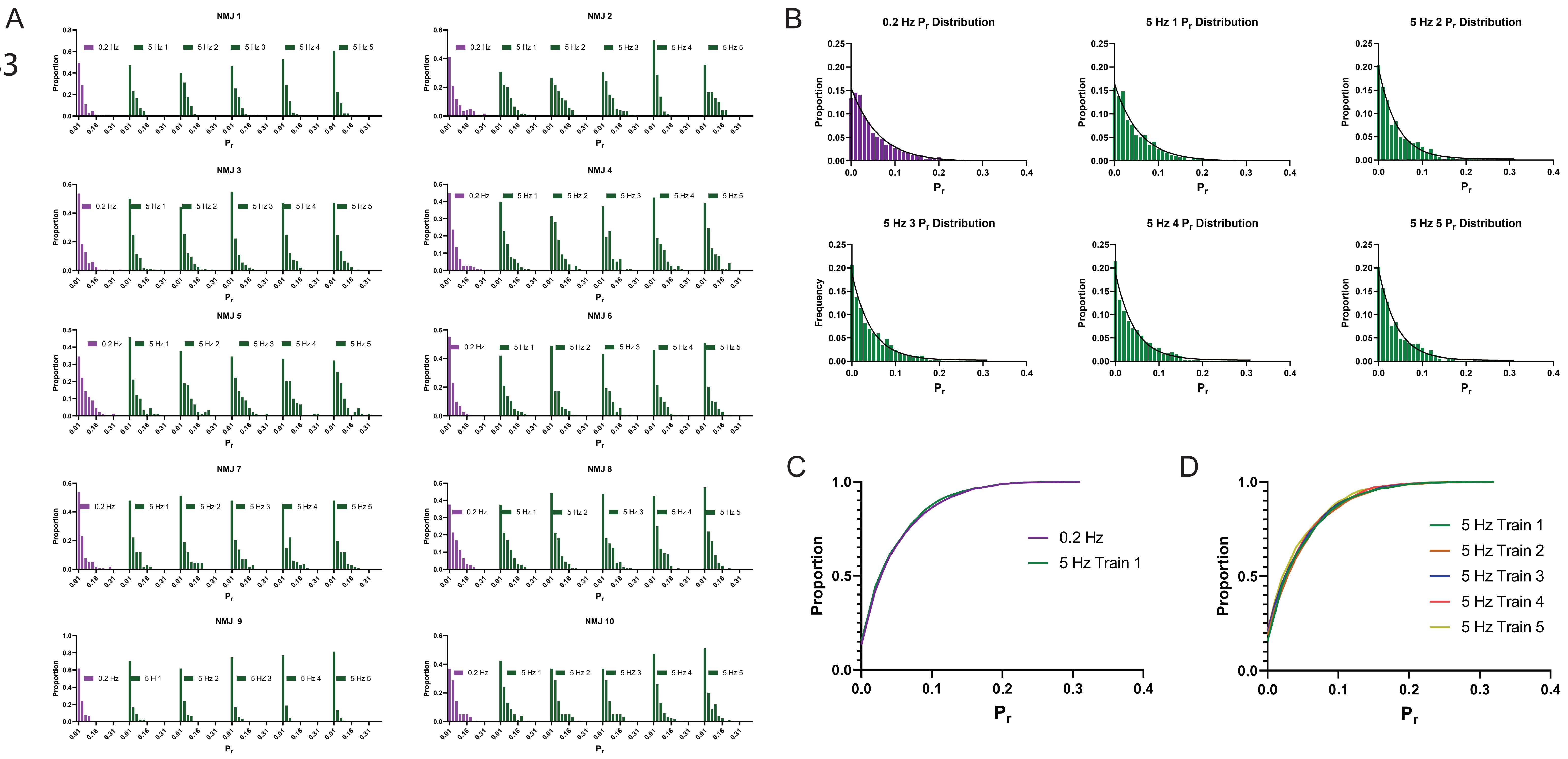
